# Platelet Activating Factor Activity Modulates Hyperoxic Neonatal Lung Injury Severity

**DOI:** 10.1101/2023.03.14.532697

**Authors:** Aaron J. Yee, Jegen Kandasamy, Namasivayam Ambalavanan, Changchun Ren, Brian Halloran, Nelida Olave, Teodora Nicola, Tamas Jilling

## Abstract

Hyperoxia-induced inflammation contributes significantly to developmental lung injury and bronchopulmonary dysplasia (BPD) in preterm infants. Platelet activating factor (PAF) is known to be a major driver of inflammation in lung diseases such as asthma and pulmonary fibrosis, but its role in BPD has not been previously investigated. Therefore, to determine whether PAF signaling independently modulates neonatal hyperoxic lung injury and BPD pathogenesis, lung structure was assessed in 14 day-old C57BL/6 wild-type (WT) and PAF receptor knockout (PTAFR KO) mice that were exposed to 21% (normoxia) or 85% O_2_ (hyperoxia) from postnatal day 4. Lung morphometry showed that PTAFR KO mice had attenuated hyperoxia-induced alveolar simplification when compared to WT mice. Functional analysis of gene expression data from hyperoxia-exposed vs. normoxia-exposed lungs of WT and PTAFR KO showed that the most upregulated pathways were the *hypercytokinemia/hyperchemokinemia* pathway in WT mice, *NAD signaling* pathway in PTAFR KO mice, and *agranulocyte adhesion and diapedesis* as well as other pro-fibrotic pathways such as *tumor microenvironment* and *oncostatin-M signaling* in both mice strains, indicating that PAF signaling may contribute to inflammation but may not be a significant mediator of fibrotic processes during hyperoxic neonatal lung injury. Gene expression analysis also indicated increased expression of pro-inflammatory genes such as CXCL1, CCL2 and IL-6 in the lungs of hyperoxia-exposed WT mice and metabolic regulators such as HMGCS2 and SIRT3 in the lungs of PTAFR KO mice, suggesting that PAF signaling may modulate BPD risk through changes in pulmonary inflammation and/or metabolic reprogramming in preterm infants.

## INTRODUCTION

Bronchopulmonary dysplasia (BPD) is a disorder characterized by abnormal lung development and vascular remodeling that contributes significantly to morbidity and mortality in very preterm infants (1). Inflammatory injury caused by reactive oxygen species (ROS) generated as a result of prolonged hyperoxia exposure is a major pathophysiologic contributor to BPD (2–4). While hyperoxia-induced inflammation is often initiated by resident macrophages and other lung cells, further injury and disrupted development of the lung is induced by migrating leukocytes such as neutrophils and macrophages which release potent inflammatory mediators upon recruitment to the lungs (5–7).

Platelet activating factor (PAF, 1-alkyl-2-acetyl-*sn*-glycero-3-phosphocholine) is one such inflammatory mediator released by several cell types after production via cytoplasmic phospholipase A2 (PLA2) isoforms, and/or produced extracellularly via secretory isoforms of PLA2. PAF activity is mediated through its cognate receptor (PTAFR), upon binding to which it activates production of lipid mediators via activation of cyclooxygenases and lipoxygenases as well as transcriptional activation of inflammatory cytokines such as IL-8 in humans and its murine equivalent CXCL1 (GRO/KC) that induce further influx of activated neutrophils and macrophages to the lung (8–10). PAF is also known to directly decrease gas exchange in the lung by increasing airway reactivity and pulmonary vascular permeability and has been implicated as a mediator in the pathogenesis of pulmonary disorders such as acute respiratory distress syndrome (ARDS) and asthma (11–13).

Increased PAF levels in bronchoalveolar lavage (BAL) fluid and serum are known to be associated with higher risk for severe BPD in preterm infants and in animal models (14, 15). However, no previous studies have evaluated whether PAF signaling independently modulates neonatal hyperoxic lung injury and BPD pathogenesis. Therefore, in the current study, we hypothesized that PAF signaling through PTAFR is a key regulator of hyperoxia-induced lung inflammation and injury that contributes to BPD pathogenesis. We tested our hypothesis by first determining whether hyperoxia causes changes in PAF signaling mediators such as PTAFR and isoforms of enzymes such as phospholipase A2 G2E (PLA2G2E) and PAF acetyl hydrolase (PLA2G7) that modulate PAF metabolism (**Figure 1A**) in the lungs (16, 17). We then evaluated newborn PTAFR gene-targeted (PTAFR KO) mice with decreased PTAFR expression in our well-established hyperoxia-induced BPD model and measured changes caused by downregulation of PAF signaling in lung structure and function, gene expression and cytokine content in the lungs of these mice compared to C57BL6 WT mice.

**Figure 1:**
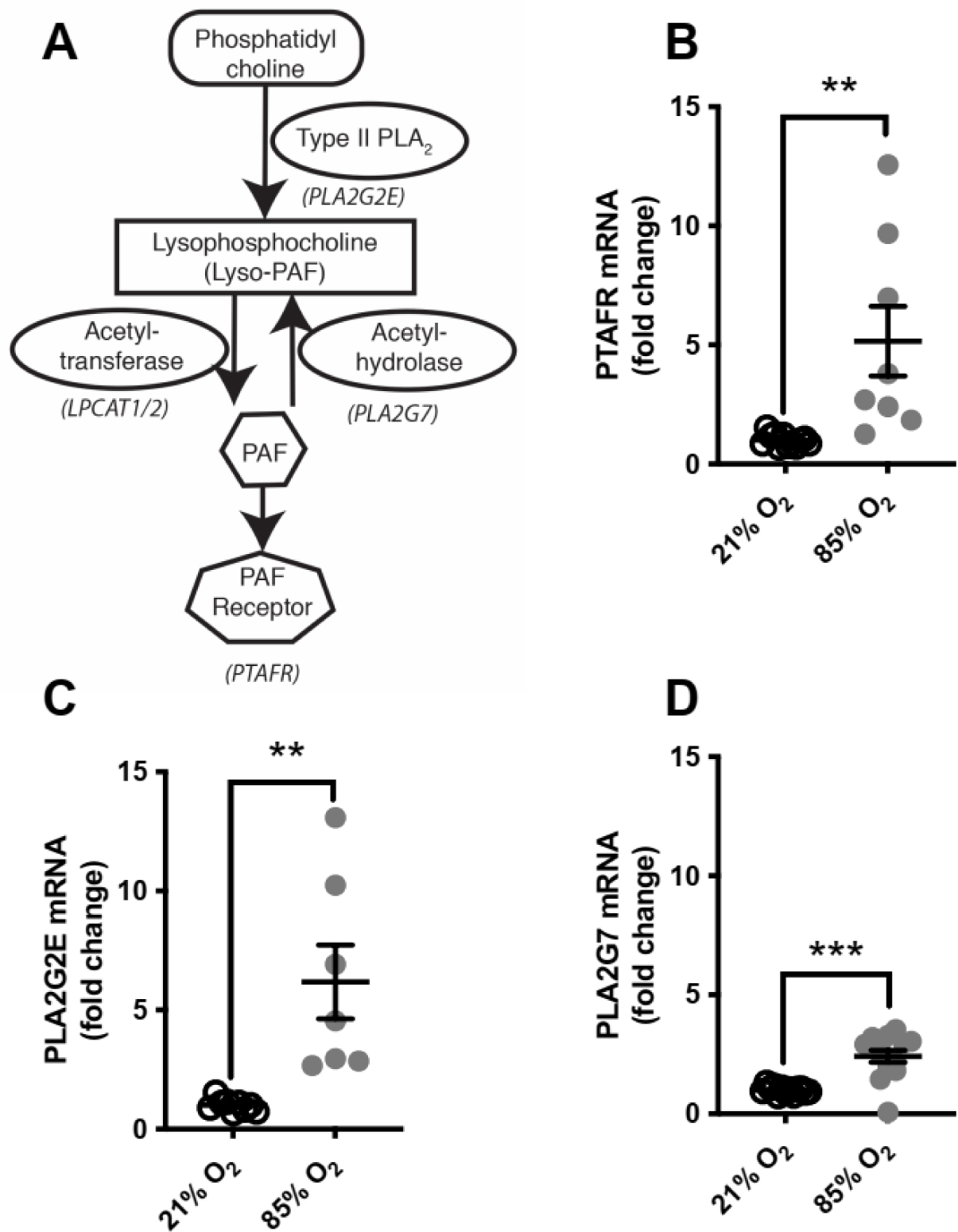
Gene expression changes of PAF-related genes. (**A**) Schematic of PAF homeostasis. Upon inflammatory activation, PAF is synthetized by type II PLA2 and lysophosphatidylcholine acyltransferase. Lysophosphatidylcholine (Lyso-PAF) is an intermediate with limited biological activity and it is converted to PAF by transfer of an acetyl group in the sn-2 position. Type 7 PLA2 (PAF acetylhydrolase) mediates the inactivation of PAF by removal of the acetyl group. Gene expression of key regulators of PAF activity such as (**B**) Platelet activating factor receptor (PTAFR) (**C**) Phospholipase A2G2E (PLA2G2E) and (**D**) PAF acetyl hydrolase (PLA2G7) were quantified by real time PCR using RNA extracted from lungs of mice exposed to normoxia or hyperoxia. ** P<0.01, *** P<0.001; t test.

## METHODS

### BPD model

The Institutional Animal Care and Use Committee of the University of Alabama at Birmingham approved all protocols. C57BL/6 (wild-type or WT) mice purchased from Charles River and C57BL/6 mice with “loss of function” of the PTAFR gene without PAF receptor expression (PTAFR KO; decreased PAF signaling) kindly provided by Dr. Shimizu were used to generate timed-pregnant mice by mating of females with single-housed male mice (8). Newborn mouse littermates and their dams were exposed to either normoxia (21% O_2_; control group) or hyperoxia (85% O_2;_ experimental group) from postnatal day 4 (P4) to P14 to induce lung injury as previously described (18).

### Lung Function

Hyperoxia or normoxia exposed P14 mice were anesthetized using isoflurane to measure lung compliance using a FlexiVent apparatus (SCIREQ, Montreal, QC, Canada) as previously described (19, 20).

### Lung structure

Alveolar morphometry was performed by quantifying using mean linear intercept (MLI) and radial alveolar counts (RAC) in 5 µM H&E stained sections of lungs from P14 mice using a TE2000U microscope (Nikon), QiCam Fast Cooled high-resolution CCD camera and MetaMorph 7.10 (Universal Imaging) as previously described (19, 20).

### Gene expression and pathway analysis

Whole lung RNA samples (with RNA integrity > 8) were submitted to Novogene (www.novogene.com) where library preparation and sequencing was performed according to Illumina protocols. Raw read counts received from Novogene were analyzed using the public access Galaxy server and a fastp-RNA STAR-featurecount-DEseq2 pipeline. Ingenuity pathway analysis (IPA; Ingenuity Systems, CA) was used to identify canonical pathways, genes and networks that were differentially regulated by hyperoxia exposure. A fold change cutoff of ≥ ±2 and false discovery rate (q) <0.05 was used to generate and control for the occurrence of false discoveries in downstream data sets. Submission of the microarray data to GEO is in progress.

### Quantitative PCR (qPCR)

Total RNA was isolated from lung homogenates using a Qiagen RNEasy Mini kit, quantified using Quant-iT RNA RiboGreen assay kit (Thermo Fisher), and reverse transcribed using PrimeScript RT mastermix (Takara). Quantitative real-time PCR (qPCR) was performed using a Rotor-Gene Q instrument (Qiagen). TaqMan MGB primer/probe sets and Eukaryotic 18S rRNA VIC primer/probe set endogenous control (Thermo Fisher) with Premix Ex Taq master mix (Takara) were used to analyze genes of interest including the chemokine *CXCL1* (Mm04207460_m1), and mediators of NAD signaling such as *HMGCS2* (Mm00550050_m1) and *SIRT3* (Mm00452131_m1).

### BAL cytokines

BAL samples were obtained by lavaging lungs with 0.3 ml of PBS intratracheally at least twice, flash-frozen in dry ice, and stored for protein analysis. BAL fluid was diluted two-fold and analyzed using the V-PLEX Proinflammatory Panel 1 (mouse) kit and analyzed using the MESO Quickplex SQ 120 (MSD Technologies).

### Statistical methods

Data were expressed as means ± SEM. Two-way ANOVA was used to test for separate and combined effects of PAF and hyperoxia using gene-targeted mice. Multiple-comparison testing (Tukey) was performed if overall statistical significance was *P<*0.05. GraphPad Prism version 7.00 for Mac OS X (GraphPad Software, La Jolla California USA, www.graphpad.com.) was used for all data analysis.

## RESULTS

### Hyperoxia exposure increases expression of PAF activity regulators in the newborn lung

Hyperoxia exposure increased mRNA expression of *PTAFR* (5.2 ± 1.5-fold; p>0.01) in lung homogenates of newborn mice. Hyperoxia also increased expression of *PLA2G2E* (which is a type II, secretory PLA2 isoform) in lung homogenates of WT mice (6.2 ± 1.6-fold; p>0.01). Additionally, PLA2G7, an isoform of PAF-AH known to be active in lung tissue, had increased expression (**Figures 1B-D**) in response to hyperoxia in these mice, but to a lower degree than either *PTAFR* or *PLA2G2E* (2.4±0.3-fold; P<0.001).

### Absence of PTAFR decreases hyperoxia-induced alveolar simplification

Lungs of normoxia-exposed PTAFR KO mice (**Figure 2A**) had similar sized alveoli and degree of septation when compared to lungs of normoxia-exposed WT mice (**Figure 2B**). When exposed to hyperoxia, lung sections of hyperoxia-exposed PTAFR KO mice (**Figure 2C**) mice showed improved alveolarization (as shown by the smaller alveolar sizes and increased septation) compared to WT controls (**Figure 2D**). These changes in alveolar size and septation were also quantified through morphometric analysis of images. Mean linear intercept (MLI), which is an indication of alveolar size, was increased in hyperoxia-exposed PTAFR and WT mice compared to their normoxia-exposed controls. However, this hyperoxia-induced pathologic increase in alveolar size was attenuated in hyperoxia-exposed PTAFR KO mice compared to WT mice (**Figure 2E**). Radial alveolar count (RAC), which reflects the magnitude of alveolar septation, was decreased in hyperoxia-exposed PTAFR KO and WT mice compared to their normoxia-exposed controls. Similar to the changes noted in MLI, this hyperoxia-induced pathologic decrease in alveolar count was also attenuated in hyperoxia-exposed PTAFR KO mice when compared to their WT mice counterparts (**Figure 2F**).

**Figure 2:**
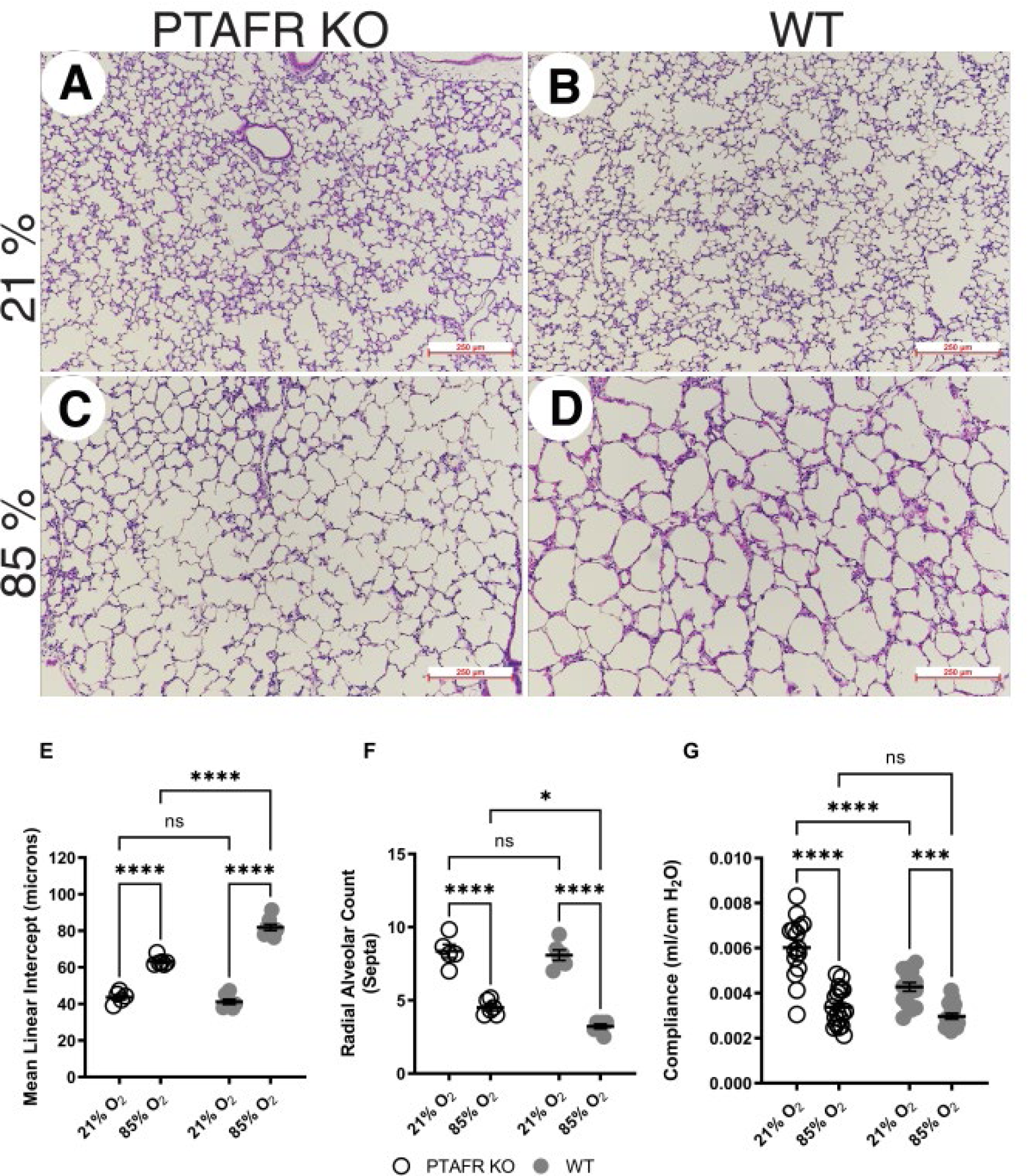
Effects of PAF signaling downregulation on hyperoxia-induced lung injury. (**A**-**D**) Representative images from PAF receptor gene targeted mice (PTAFR KO) and wild type (WT) mice (left to right) exposed to normoxia (top) or hyperoxia (bottom). 3 images collected from 20 histological sections collected from 6 mice per group were used to quantify mean linear intercept (**E**) and radial alveolar count (**F**). Lung compliance (**G**) was assessed using FlexiVent. ** P<0.01, *** P<0.001, **** P<0.0001, ns – not significant; two-way ANOVA and Tukey test for post hoc analysis.

### The absence of PTAFR does not affect hyperoxia-induced reduction in lung compliance

Hyperoxia-induced lung injury is associated with decreased lung compliance and, as would be expected, the hyperoxia-induced alveolar simplification noted in PTAFR KO and WT mice was also accompanied by reduction in lung compliance when compared to their normoxia-exposed controls. Lung compliance was also lower in normoxia exposed WT mice compared to PTAFR KO mice. However, and contrary to expectations based on the changes in alveolar size noted in hyperoxia-exposed PTAFR KO vs. WT mice, no differences were noted in lung compliance between hyperoxia-exposed PTAFR KO vs. WT mice (**Figure 2G**).

### The absence of PTAFR modifies hyperoxia-induced gene expression in the newborn lung

RNA-seq analysis was performed to analyze gene expression in lung homogenates from normoxia and hyperoxia exposed PTAFR KO and WT newborn mice. Substantial separation was noted between the lung transcriptome clusters of hyperoxia-exposed PTAFR KO mice (red) and WT mice (green) on principal component analysis (PCA) but not between the lung transcriptomes of normoxia-exposed PTAFR KO mice (blue) and WT (purple, indicating that gene expression in the lungs was differentially modified by hyperoxia exposure between these 2 groups of mice (**Figure 3A**).

**Figure 3:**
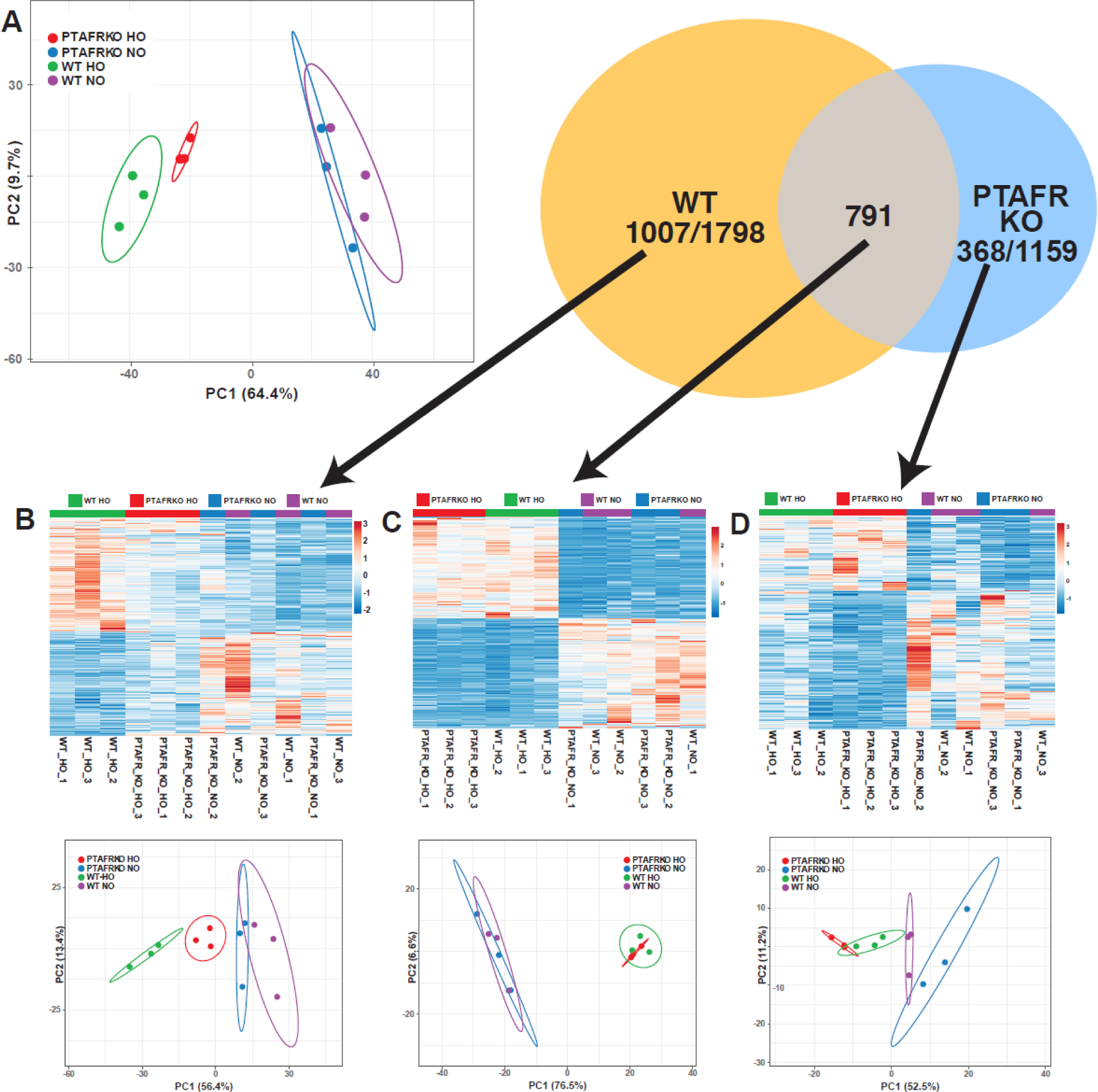
Effects of PAF signaling downregulation on hyperoxia-induced gene expression. (**A**) Principal component analysis (PCA) of all differentially expressed genes in the lungs of WT mice and PTAFR KO mice indicates overlap between normoxia exposed samples and clear separation between hyperoxia exposed WT mice and PTAFR KO mice. (**B-D**) Heat maps and PCA plots of the 1007 genes found to be differentially expressed in WT mice only (**B**), the 791 genes differentially expressed in both WT and PTAFR KO mice (**C**) and the 368 genes differentially expressed genes in PTAFR KO mice only (**D**).

Further analysis found that hyperoxia exposure caused 1798 transcripts to be differentially regulated (defined as TPM>0.1, fold change > ±2 and q <0.05) in the lungs of WT mice, of which 1007 were regulated by hyperoxia in WT only. Robust separation was noted between the hyperoxia-exposed PTAFR KO (red) and WT (green) mice on both the PC plot and the heatmap of the transcripts in the lung that were differentially regulated by hyperoxia in WT only, while the normoxia exposed WT (purple) and PTAFR KO (blue) mice exhibited significant overlap. Similarly, hyperoxia exposure caused 1159 transcripts to be differentially expressed in the lungs of PTAFR KO mice of which 368 were found to be regulated by hyperoxia in PTAFR KO mice only. The separation of hyperoxia exposed PTAFR KO and WT groups is less robust on PC plot, but still present in heatmap hierarchical clustering generated from transcript counts of genes regulated in PTAFR KO only. Notably, the separation of normoxia exposed PTAFR KO and WT groups is increased in this gene set (**Figure 3B-D**). 791 transcripts were found to be differentially regulated by hyperoxia in the lungs of both WT and PTAFR KO mice. Hyperoxia-exposed PTAFR KO and WT groups are not separated by hyperoxia on PC plot but still exhibit hierarchical clustering on heat map generated from transcript counts of genes that are regulated in both genotypes by hyperoxia (**Figure 3C**).

**Supplemental Tables 1 and 2** show the top 20 genes that were upregulated or downregulated by hyperoxia exposure in these mice, along with the corresponding fold change and FDR p-values for the same genes from the PTAFR KO hyperoxia vs normoxia comparison. Genes with > 2-fold difference in the fold change in the WT comparison and FDR p < 0.05 in both comparisons are highlighted in yellow in these tables. Known functional genes highlighted in **Supplemental Table 1** included Ccl2, Cxcl10, Cxcl1 (which are involved in monocyte and neutrophil chemotaxis) and Dio1 (involved in thyroid hormone metabolism). Known functional genes highlighted in **Supplemental Table 2** included Wfdc8, Wfdc6a (peptidase inhibitors), Ear1(eosinophil chemotaxis), and Abca17 as well as Fabp1(involved in lipid metabolism). **Supplemental Tables 3 and 4** show the top 20 genes that were upregulated or downregulated by hyperoxia in these mice, along with the corresponding FDR and p-values for these genes from the WT hyperoxia vs. normoxia comparison. Genes with > 2-fold difference in the fold change in the PTAFR KO comparison and FDR p < 0.05 in both comparisons are highlighted in yellow in these tables. Known functional genes highlighted in **Supplemental Table 3** included Mcpt4, Cma1 and Tpsb2 (involved in mast cell function), Aldh1a3 (associated with metabolic regulation in pulmonary arterial smooth muscle cells), Pla1a (a phospholipase) and Shisa6 (known to be involved in cellular differentiation). Known functional genes highlighted in **Supplemental Table 4** included Kcgn4 (potassium channel regulator), Cpa1 (a carboxypeptidase), Cyp2e1 (known to be associated with xenobiotic metabolism and lung development), Cbln1 (known to be associated with pulmonary hypertension, Crabp1 (involved in retinoic acid metabolism) and Wfdc8.

### The absence of PTAFR modifies hyperoxia-induced effects on pathways in the neonatal lung

IPA analysis was performed to categorize cell-specific pathways, functions and molecules previously known to be involved in regulating functions of lung epithelial and endothelial cells, fibroblasts, smooth muscle cells, immune cells, and mesenchymal stem cells that were differentially enriched using the 1007 genes differentially regulated only in WT mice, the 368 genes differentially regulated only in PTAFR KO mice as well as the 791 genes differentially regulated in both WT and PTAFR KO mice. Processes and molecules involved in immune cell recruitment and migration were found to be differentially upregulated by hyperoxia in the lungs of WT mice, whereas processes and molecules related to activation of immune cells were downregulated in the lungs of PTAFR KO mice. Processes and molecules related to lymphocyte, neutrophil and macrophage function and movement were found to upregulated by hyperoxia in the lungs of both groups (**Figure 4**). Canonical pathways that were found to be differentially regulated with absolute activation Z scores ≥ 2 and P <0.05 (-log values > 1.3) from these analyses are listed in **Table 1**. *Hypercytokinemia/hyperchemokinemia in the Pathogenesis of Influenza pathway*, *Nod1/Nod2 signaling pathway* and *IL-33 signaling pathway*, which were the top canonical pathways differentially upregulated by hyperoxia in the lungs of WT mice only and not in the lungs of PTAFR KO mice (and therefore considered to be PTAFR-dependent genes and pathways in this analysis), are all known to be involved in innate immune response, inflammation, chemokine and cytokine production. An IPA generated pathway map of the *hypercytokinemia/hyperchemokinemia* pathway is shown in **Figure 5**. Fold changes and corresponding p-values for the expression of individual genes from these pathways are shown in **Tables 2, 3,** and **4**. The top pathway differentially enriched by hyperoxia in the lungs of PTAFR KO mice but not in the lungs of WT mice was the *NAD signaling pathway* which is involved in cellular metabolic, antioxidant and immune responses. An IPA generated pathway map of the *NAD signaling pathway* is shown in **Supplemental Figure 1**. Fold changes and corresponding p-values for the expression of individual genes from this pathway are shown in **Table 5** and indicate that the mitochondrial enzyme *HMGCS2* is the most differentially upregulated protein in the PTAFR KO mice lung compared to the WT mice lung. This finding was validated by RT-PCR analysis of HMGCS2 mRNA in the lungs of these mice. Additionally, and though this was not noted in the RNA-seq and IPA studies, expression of SIRT3 (a major regulator of HMGCS2) was also noted to be regulated by hyperoxia exposure and higher in PTAFR KO mice lungs compared to WT mice lungs (**Supplemental Figure 2**). Pathways that were noted to be differentially regulated by hyperoxia in both groups included *Agranulocyte adhesion and diapedesis* (**Supplemental Table 5**), *pathogen induced cytokine storm signaling* and several pathways associated with fibrotic processes such as the *tumor microenvironment* (**Supplemental Table 6**), *oncostatin M signaling* and *wound healing* pathways. An IPA generated pathway map of the *tumor microenvironment pathway* is shown in **Supplemental Figure 3**.

**Figure 4:**
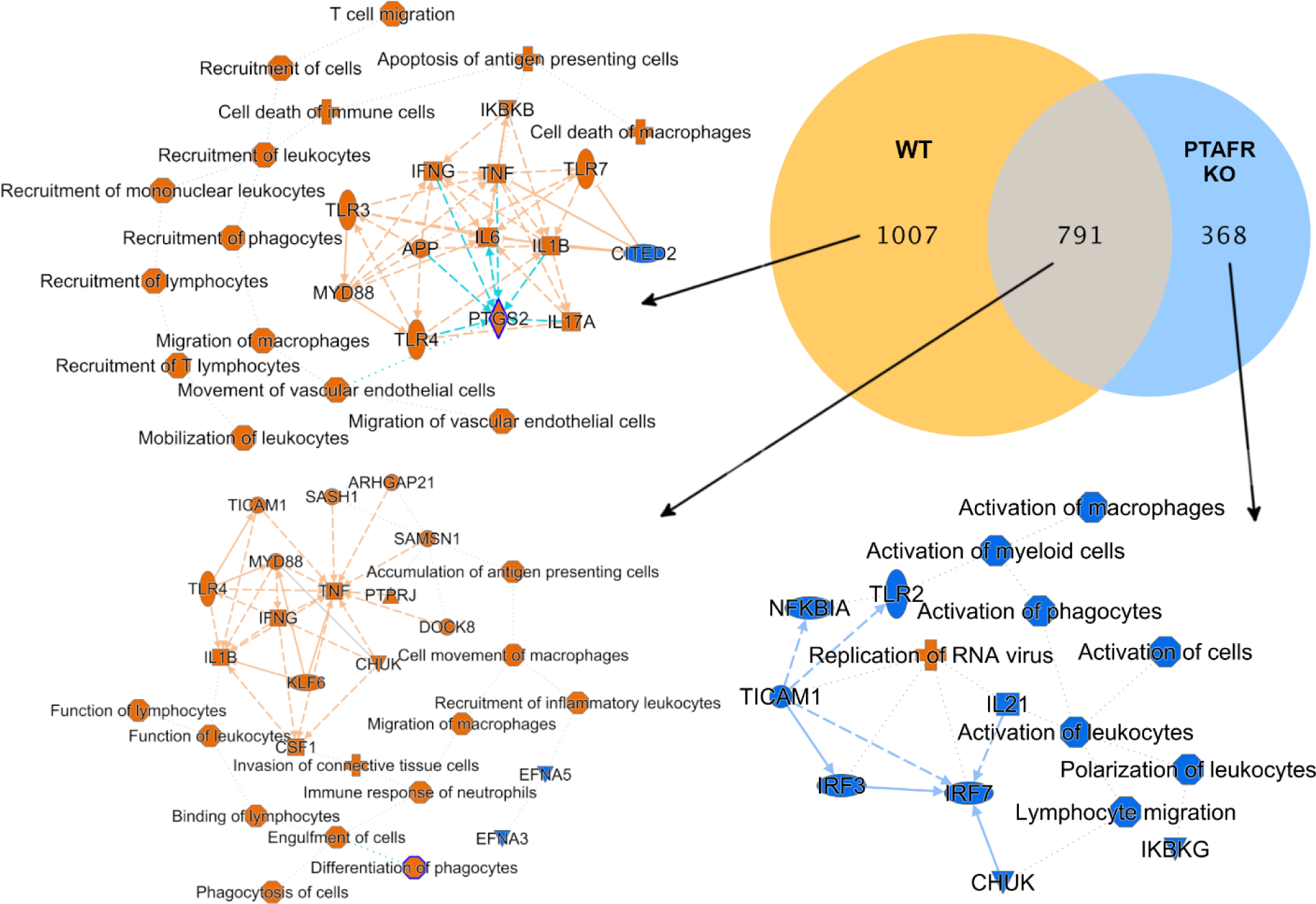
Network analysis of lung and immune cell function related genes that were differentially expressed in hyperoxia-exposed lungs. Genes known to be associated with lung and immune cell function that were differentially upregulated (fold changes > 2 and FDR P-value < 0.05) by hyperoxia in the lungs of mice exposed to hyperoxia compared to mice exposed to normoxia were analyzed using Ingenuity IPA to identify molecules, processes, diseases, and pathways that were differentially regulated (orange for upregulated and blue for downregulated) by hyperoxia in the lungs of normoxia-exposed lungs vs. hyperoxia-exposed lungs.

**Figure 5:**
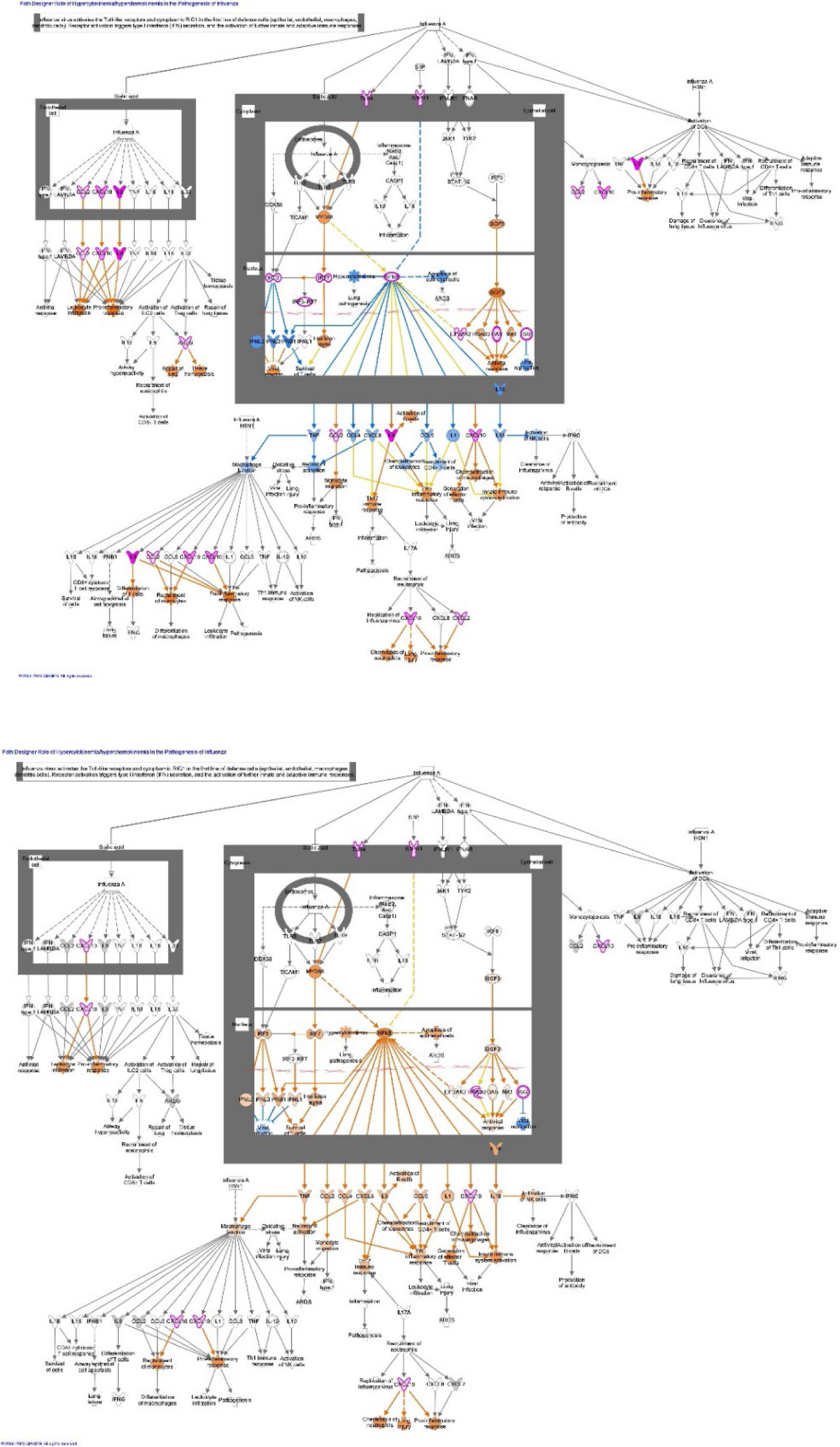
Pathway map of the *hypercytokinemia/hyperchemokinemia* pathway. Known interactions and interrelationships between the component genes of the *hypercytokinemia/hyperchemokinemia* pathway that was identified to be differentially regulated by hyperoxia in the lungs of WT mice only and not in the lungs of PTAFR KO mice.

**Table 1:** Canonical pathways related to lung and immune cell function that were regulated with Z(abs) ≥ 2 and −logP >1.3 (P<0.05) by hyperoxia in the lungs of WT only, PTAFR KO only and in both.

| Group | Canonical Pathway | $-\log P$ | Ratio | Z-score |
| --- | --- | --- | --- | --- |
| WT | Role of Hypercytokinemia/hyperchemokine in the Pathogenesis of Influenza | 2.43 | 0.0921 | 2.646 |
|  | Nod1/Nod2 Signaling Pathway | 1.38 | 0.0511 | 2.333 |
|  | IL-33 Signaling Pathway | 1.36 | 0.0508 | 2.333 |
|  | Nicotine Degradation II | 1.79 | 0.0877 | 2.236 |
|  | Tumor Microenvironment Pathway | 2.18 | 0.0625 | 2.111 |
|  | Estrogen Biosynthesis | 1.6 | 0.093 | 2 |
|  | Nicotine Degradation III | 1.33 | 0.0769 | 2 |
|  | IL-17 Signaling | 3.32 | 0.0774 | 1.387 |
|  | UVA-Induced MAPK Signaling | 1.87 | 0.0722 | -2 |
| Both | Agranulocyte Adhesion and Diapedesis | 7.68 | 0.112 | 3.342 |
|  | Pathogen Induced Cytokine Storm Signaling Pathway | 4.99 | 0.0734 | 3.138 |
|  | Tumor Microenvironment Pathway | 3.82 | 0.0852 | 2.324 |
|  | Role Of Chondrocytes in Rheumatoid Arthritis Signaling Pathway | 3.22 | 0.0863 | 2.309 |
|  | Oncostatin M Signaling | 2.15 | 0.116 | 2.236 |
|  | Osteoarthritis Pathway | 3.02 | 0.0693 | 2.138 |
|  | Wound Healing Signaling Pathway | 4.19 | 0.0785 | 2.065 |
| PTAFR-KO | NAD Signaling Pathway | 2.01 | 0.035 | 2.236 |
|  | IL-13 Signaling Pathway | 2.63 | 0.049 | -2.236 |

**Table 2:** Genes included in the *Role of Hypercytokinemia/hyperchemokinemia in the Pathogenesis of Influenza* IPA canonical pathway. Genes that were significantly regulated by hyperoxia in the lungs of WT and PTAFR KO mice (fold change values > 2 vs. the other group and P < 0.05 in both groups) are highlighted in yellow.

| Gene | WT HO vs. WT NO |  | PRKO HO vs. PRKO NO |  |
| --- | --- | --- | --- | --- |
|  | Fold change | FDR p-value | Fold change | FDR p-value |
| **IL6** | 59.13956 | 0.000334 | 18.35983 | 0.000402 |
| **Ccl2** | 39.48902 | 1.22E-17 | 13.99874 | 8.82E-19 |
| **Cxcl10** | 22.33504 | 2.09E-18 | 4.326959 | 8.97E-06 |
| **Cxcl1** | 22.16526 | 1.79E-07 | 2.707381 | 1.19E-03 |
| **Cxcl2** | 9.263299 | 0.000483 | 1.953732 | 0.506267 |
| **Tnf** | 4.456041 | 0.021756 | 1.602264 | 0.583136 |
| **Areg** | 2.780736 | 0.000883 | 1.825641 | 0.091329 |
| **Irf7** | 2.540168 | 6.97E-06 | 1.20622 | 0.590221 |
| **Tlr4** | 1.88965 | 0.000115 | 1.531674 | 0.023913 |
| **S1pr1** | 1.780886 | 0.000245 | 1.759765 | 0.000486 |
| **Tlr7** | 1.770118 | 0.104119 | 1.386386 | 0.475608 |
| **Eif2ak2** | 1.62513 | 0.008741 | 1.2871 | 0.300794 |
| **Il1b** | 1.601683 | 0.325778 | -1.59882 | 0.419898 |
| **Myd88** | 1.362539 | 0.060493 | 1.30703 | 0.155025 |
| **Irf9** | 1.321219 | 0.090967 | 1.020576 | 0.951371 |
| **Ddx58** | 1.308314 | 0.129633 | -1.03132 | 0.929511 |
| **Ifnlr1** | 1.273678 | 0.375108 | 1.479787 | 0.151646 |
| **Tlr9** | 1.102537 | 0.882486 | 1.034647 | 0.972536 |
| **Jak1** | 1.072763 | 0.743939 | -1.07194 | 0.805401 |
| **Tlr3** | 1.070181 | 0.826956 | -1.10568 | 0.792739 |
| **Casp1** | 1.014873 | 0.981614 | -1.02829 | 0.974193 |
| **Il33** | -1.09659 | 0.72618 | -1.06217 | 0.869243 |
| **Rsad2** | -1.1779 | 0.62971 | -2.81836 | 1.3E-05 |
| **Ticam1** | -1.25589 | 0.302469 | -1.04418 | 0.912393 |
| **Tyk2** | -1.41092 | 0.053697 | -1.26515 | 0.28572 |
| **Il18** | -1.54298 | 0.226694 | -1.1429 | 0.81616 |
| **Irf3** | -1.71369 | 0.000174 | -1.27283 | 0.201418 |
| **Ccl5** | -1.84365 | 0.124739 | -2.42047 | 0.024535 |

**Table 3:** Genes included in the *Nod1/Nod2 Signaling* IPA canonical pathway. Genes that were significantly regulated by hyperoxia in the lungs of WT and PTAFR KO mice (fold change values 2 vs. the other group and P < 0.05 in both groups) are highlighted in yellow.

| Gene | WT HO vs. WT NO |  | PRKO HO vs. PRKO NO |  |
| --- | --- | --- | --- | --- |
|  | Fold change | FDR p-value | Fold change | FDR p-value |
| **IL6** | 59.13956 | 0.000334 | 18.35983 | 0.000402 |
| **Ccl2** | 39.48902 | 1.22E-17 | 13.99874 | 8.82E-19 |
| **Tnfaip3** | 1.924544 | 3.91E-05 | 1.658293 | 0.003284 |
| **Map3k8** | 2.569409 | 0.001489 | 1.559903 | 0.256431 |
| **Traf3** | 1.50067 | 0.007278 | 1.341609 | 0.097254 |
| **Irf7** | 2.540168 | 6.97E-06 | 1.20622 | 0.590221 |
| **Traf6** | 1.378393 | 0.049712 | 1.061951 | 0.839143 |
| **Irf3** | -1.71369 | 0.000174 | -1.27283 | 0.201418 |
| **Trim27** | -1.43575 | 0.01888 | -1.29342 | 0.162081 |
| **Slc15a4** | -1.41817 | 0.04805 | -1.44923 | 0.045364 |
| **Nod1** | -1.46643 | 0.02373 | -1.56845 | 0.008421 |
| **Card9** | -4.46054 | 8.33E-14 | -3.34982 | 2.29E-09 |

**Table 4:** Genes included in the *IL33* signaling IPA canonical pathway. Genes that were significantly upregulated by hyperoxia in the lungs of WT and PTAFR KO mice (fold change values 2 vs. the other group and P < 0.05 in both groups) are highlighted in yellow (WT higher than PTAFR KO) or blue (PTAFR KO higher than WT).

| Gene | WT HO vs. WT NO |  | PRKO HO vs. PRKO NO |  |
| --- | --- | --- | --- | --- |
|  | Fold change | FDR p-value | Fold change | FDR p-value |
| **IL6** | 59.13956 | 0.000334 | 18.35983 | 0.000402 |
| **Ccl2** | 39.48902 | 1.22E-17 | 13.99874 | 8.82E-19 |
| **Mcpt4** | 5.902817 | 6.04E-08 | 25.13168 | 2.89E-24 |
| **Cma1** | 4.695365 | 4.02E-07 | 19.51972 | 4.14E-25 |
| **Mmp12** | 4.596144 | 4.3E-05 | 1.778731 | 0.262224 |
| **Ptgs2** | 4.301681 | 2.07E-06 | 1.871583 | 0.103458 |
| **Tpsb2** | 4.100237 | 2.16E-07 | 17.83594 | 1.15E-29 |
| **Ccl17** | 3.592802 | 0.000583 | 1.526953 | 0.434723 |
| **Casp4** | 3.303143 | 5.75E-05 | 2.086256 | 0.030832 |
| **Areg** | 2.780736 | 0.000883 | 1.825641 | 0.091329 |
| **Icam1** | 2.670234 | 2.38E-06 | 2.442992 | 3.05E-05 |
| **Map3k8** | 2.569409 | 0.001489 | 1.559903 | 0.256431 |
| **Mmp2** | 1.965964 | 7.86E-05 | 2.159559 | 5.68E-06 |
| **Prkar2a** | 1.622395 | 0.004136 | 1.378 | 0.11513 |
| **Vcam1** | 1.567566 | 0.004243 | 1.569551 | 0.005855 |
| **Nfkb1** | -1.55138 | 0.002149 | -1.48753 | 0.009442 |
| **Casp2** | -1.67148 | 0.000979 | -1.48012 | 0.024231 |
| **Irf3** | -1.71369 | 0.000174 | -1.27283 | 0.201418 |
| **Creb1** | -1.73869 | 0.000326 | -1.68824 | 0.001036 |
| **Casp6** | -1.7516 | 5.71E-05 | -1.43335 | 0.024982 |
| **Pik3c2b** | -1.89048 | 9.8E-05 | -1.48352 | 0.03846 |
| **Gata3** | -2.03659 | 0.003135 | -2.70205 | 2.28E-05 |
| **Pik3r6** | -2.12876 | 0.000689 | -1.48404 | 0.155649 |
| **Sigirr** | -2.34775 | 4.96E-06 | -1.56624 | 0.045108 |
| **Fbxl19** | -2.72802 | 2.74E-10 | -1.48785 | 0.045108 |
| **Kit** | -10.1834 | 3.07E-51 | -7.56701 | 2.4E-39 |

**Table 5:**
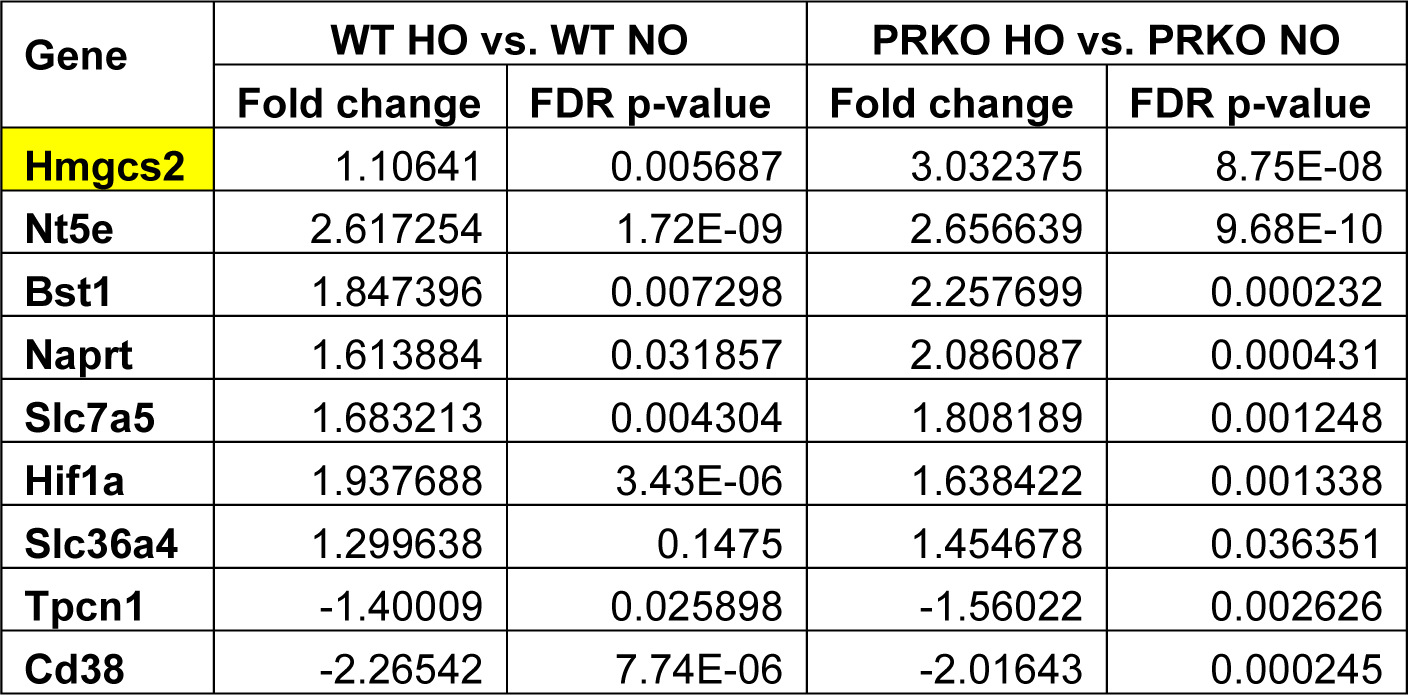
Genes included in the *NAD+ signaling* IPA canonical pathway. Genes that were significantly upregulated by hyperoxia in the lungs of WT and PTAFR KO mice (fold change values 2 vs. the other group and P < 0.05 in both groups) are highlighted in yellow.

| Gene | WT HO vs. WT NO |  | PRKO HO vs. PRKO NO |  |
| --- | --- | --- | --- | --- |
|  | Fold change | FDR p-value | Fold change | FDR p-value |
| Hmgcs2 | 1.10641 | 0.005687 | 3.032375 | 8.75E-08 |
| Nt5e | 2.617254 | 1.72E-09 | 2.656639 | 9.68E-10 |
| Bst1 | 1.847396 | 0.007298 | 2.257699 | 0.000232 |
| Naprt | 1.613884 | 0.031857 | 2.086087 | 0.000431 |
| Slc7a5 | 1.683213 | 0.004304 | 1.808189 | 0.001248 |
| Hif1a | 1.937688 | 3.43E-06 | 1.638422 | 0.001338 |
| Slc36a4 | 1.299638 | 0.1475 | 1.454678 | 0.036351 |
| Tpcn1 | -1.40009 | 0.025898 | -1.56022 | 0.002626 |
| Cd38 | -2.26542 | 7.74E-06 | -2.01643 | 0.000245 |

IPA analysis was also carried out in a non-cell specific manner using the 1007, 368 and 791 genes that were differentially regulated in the lungs of WT only, PTAFR KO only as well as both groups respectively. In addition to the upregulation of inflammatory and immune processes that was previously noted in lung and immune cell specific IPA analysis of lungs from WT mice, marked inhibition of the *LXR/RXR activation* canonical pathway that is known to modulate cholesterol and lipid metabolism was also noted in this analysis in WT mice lungs. While pathways related to immune cell activation continued to exhibit inhibition in the lungs of PTAFR KO mice, the *hepatic fibrosis /hepatic stellate cell activation pathway* which contains processes known to be involved in lung fibrosis as well was additionally identified as a pathway whose activation state could not be predicted reliably in this analysis. However, several genes that belong to this pathway were differentially expressed in both groups of mice, and this pathway was noted to be markedly activated in the IPA graphical summary defined by the gene set regulated in both WT and PTAFR KO mice indicating once again that pro-fibrotic processes were enriched by hyperoxia exposure in the lungs of both these mice groups (**Figure 6** and Supplemental Table 7).

**Figure 6:**
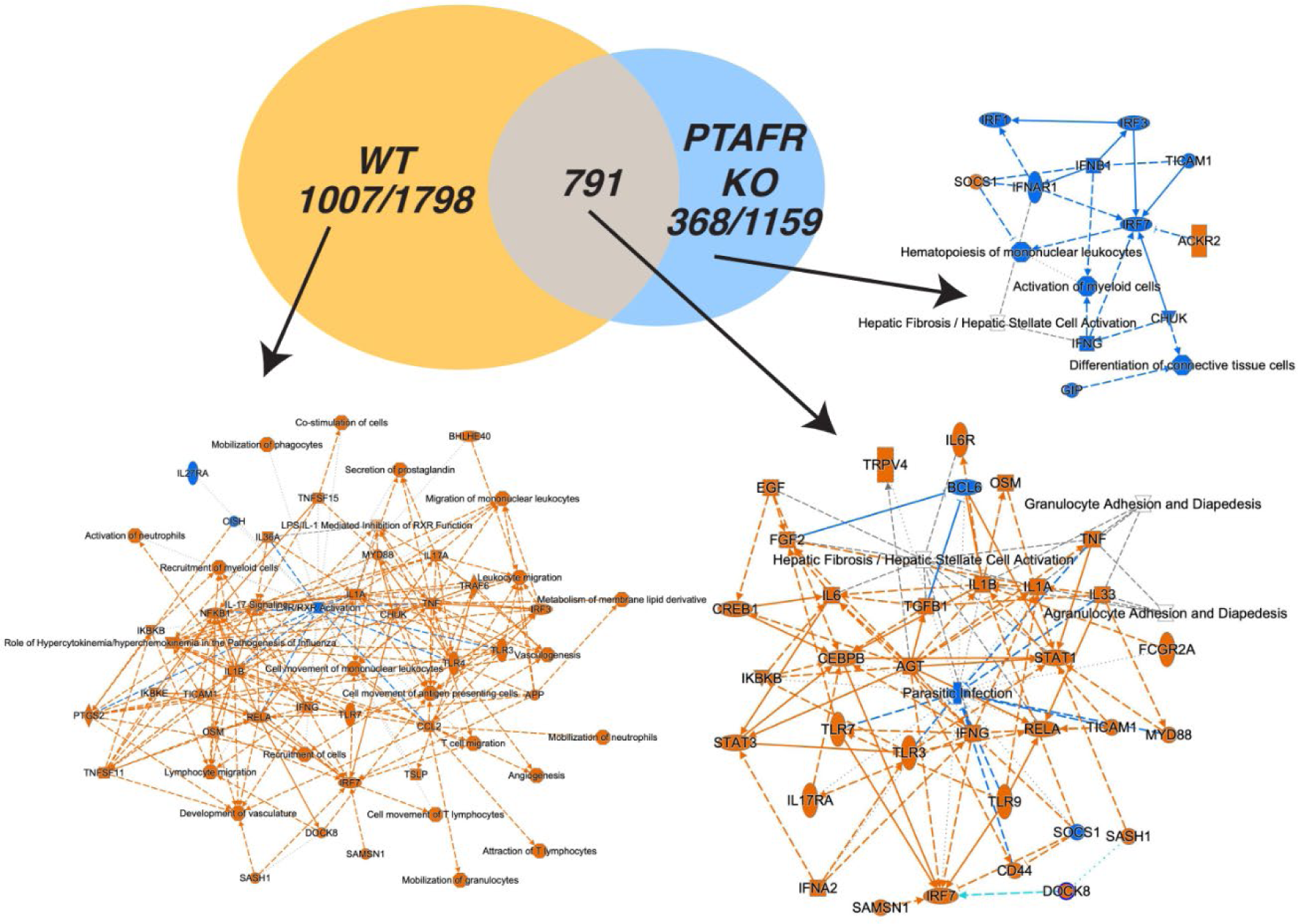
Network analysis of all genes that were differentially expressed in hyperoxia-exposed lungs. Genes that were differentially upregulated (fold changes > 2 and FDR P-value < 0.05) by hyperoxia in the lungs of mice exposed to hyperoxia compared to mice exposed to normoxia were analyzed using Ingenuity IPA to identify molecules, processes, diseases, and pathways that were differentially regulated (orange for upregulated and blue for downregulated) by hyperoxia in the lungs of normoxia-exposed lungs vs. hyperoxia-exposed lungs.

### PTAFR signaling modifies hyperoxia-induced CXCL1 expression in the developing lung

Since IPA identified activation and function of cytokines and chemokines as the top differentially activated pathway in the lungs of WT mice that had increased hyperoxic lung injury compared to PTAFR KO mice, mRNA (by RT-PCR) and protein content of CXCL1, the major neutrophil chemokine, was measured in the lungs of WT and PTAFR KO mice exposed to hyperoxia. Pulmonary CXCL1 mRNA measured by qPCR was noted to be increased by hyperoxia in both strains, but this change was markedly higher in WT mice compared to the PTAFR KO mice (**Figure 7A**). BAL fluid CXCL1 protein levels were also higher in hyperoxia-exposed WT mice compared to PTAFR KO mice (**Figure 7B**).

**Figure 7:**
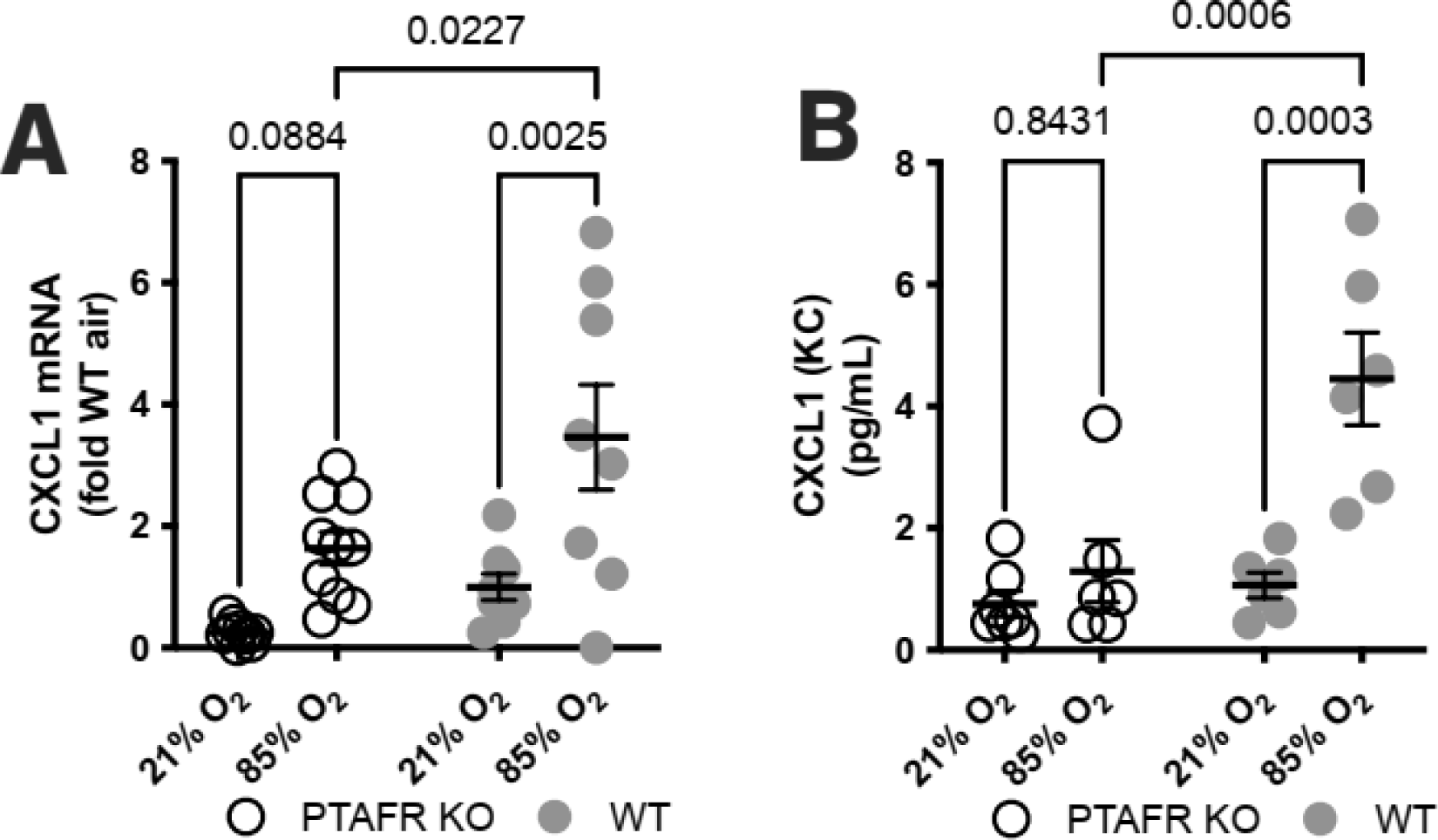
CXCL1 expression in the lung. (**A**) CXCL1 RNA measured using RNA isolated from lung tissue analyzed using real time polymerase chain reaction (RT-PCR). (**B**) CXCL1 protein content in bronchoalveolar lavage (BAL) fluid analyzed using a multiplex ELISA, two-way ANOVA and Tukey test for post hoc analysis.

## DISCUSSION

The results of this study indicate that the absence of the PAF receptor decreases hyperoxia-induced lung injury in newborn mice. Functional analysis of differentially expressed genes identified that pathways related to chemokine/cytokine activation are upregulated by a hyperoxia to a greater degree in the lungs of WT mice than in the lungs of PTAFR KO mice. Hyperoxia was also found to increase expression of enzymes in the PAF biosynthetic pathway as well as of PTAFR in the lungs of newborn mice. Overall, these findings indicate that PAF-mediated effects such as the recruitment of immune cells and the subsequent increased inflammation caused by such cells could play a significant role in hyperoxia-induced injury in the developing lung and increase risk for BPD in newborn infants.

By binding to PTAFR, its cognate receptor, PAF causes changes such as endothelial changes that can result in altered vasomotor tone, vascular permeability and chronic changes such as pulmonary vascular remodeling (10, 21–23). In addition to initiating localized inflammation on its own, PAF can produce effects such as increase in airway hyper-reactivity through the generation of eicosanoids such as thromboxane A2 and leukotrienes (24). Much of the role played by PAF in the pathogenesis of lung disorders such as chronic obstructive pulmonary disease, pulmonary fibrosis, asthma, and ARDS is mediated through its potent pro-inflammatory effects that include increased leukocyte adhesion, recruitment, chemotaxis, and degranulation. In addition, once recruited to the lung, these leukocytes in turn serve as a source of PAF themselves leading to a vicious cycle of inflammation (25, 26). PAF antagonist strategies such as recombinant human PAF-AH have been evaluated as potential therapies in animal models and human clinical trials for diseases associated with inflammation such as sepsis and necrotizing enterocolitis (27, 28). Our results suggest that such therapies may also be useful in mitigating risk for BPD in preterm infants.

Monocytes, macrophages and neutrophils have been recognized as key mediators of the inflammatory response that contributes to the hyperoxia-induced arrested alveolarization characteristic of BPD, and anti-inflammatory agents such as corticosteroids have been found to decrease risk for BPD in preterm infants (29–31). Macrophage recruitment to sites of inflammation is primarily induced by chemokines including CCL2 and macrophage-derived chemokines such as CXCL10 have been implicated in the pathobiology of lung fibrosis (32). Macrophages can be either pro-inflammatory or anti-inflammatory and the cytokine IL-6 has been known to promote alternative activation of macrophages that causes them to develop an anti-inflammatory phenotype (33). However, a recent study also indicates that macrophage abundance and alveolar injury were significantly lower in newborn IL-6^−/−^ mice exposed to hyperoxia compared to WT mice, indicating that IL-6 and macrophage-mediated mechanisms may be more pro-inflammatory than anti-inflammatory in hyperoxia-induced BPD (34). IL-6 and CCL2 also participate in Nod1/Nod2 signaling pathways, and studies indicate that Nod2 sensing of bacterial infections can lead to increased CCL2 expression and subsequent recruitment of macrophages to the lung (35). Our findings that IL-6, CCL2 and CXCL10 expression is induced to a greater degree by hyperoxia in WT mice than in PTAFR KO mice indicates that PAF activity may similarly contribute to macrophage dependent inflammation in hyperoxia-induced neonatal lung injury.

Similarly, neutrophils and IL-8 contribute to lung injury and abnormal lung development and levels of CXCL-1/IL-8 (a potent chemoattractant of neutrophils) have been found to be elevated in animal models of BPD and in tracheal aspirates of infants with BPD (3, 36, 37). Studies utilizing partial neutrophil depletion have shown reduction of tissue damage in a variety of lung injury models (38, 39). Our finding that CXCL1 was one of the most differentially regulated molecules in the lungs of WT mice exposed to hyperoxia compared to the lungs of PTAFR KO mice in this study adds to this existing evidence and suggests that decreasing IL-8 activity in the lung may be a possible mechanism through which PAF antagonists may decrease lung injury caused by BPD risk factors such as hyperoxia in preterm infants. Additionally, activation of the nuclear receptors LXR/RXR impairs neutrophil motility and reduces their influx into the lung in models of LPS-induced lung injury, suggesting that the downregulation of LXR/RXR activation pathway noted in hyperoxia-exposed WT mice (but not PTAFR KO mice) may also contribute to PAF-mediated hyperoxia-induced neonatal lung injury (40).

*IL-33 signaling pathway* was another cytokine related pathway that was found to be differentially upregulated to a greater degree by hyperoxia in WT mice compared to PTAFR KO mice in our study. IL-33 is a pleiotropic cytokine with multiple effects including activation of type 2 immune cells, promotion of inflammation and tissue remodeling. IL-33 is also known to promote lung fibrosis and disrupt alveologenesis partly through increased CCL2 and IL-6 expression in lung epithelial and endothelial cells (41–43). Previous studies have indicated that IL-33 overexpression causes pathologic changes suggestive of BPD in newborn mice (44, 45). While much of the difference in the regulation of this pathway noted between WT and PTAFR KO mice in our study was likely due to the increased IL-6 and CCL2 expression, differences were also noted in the expression of mast cell chymases such as Mcpt4, Cma1 and Tpsb2 which were found to be upregulated by hyperoxia in PTAFR KO mice compared to WT mice. Interestingly, Mcpt4, which shares homology with the human mast cell chymase Cma1, has been found to reduce airway inflammation by degrading Il-33, suggesting that upregulation of these chymases in the lungs of hyperoxia-exposed PTAFR KO mice may also contribute to decreased lung injury noted in these mice by reducing IL-33 pro-inflammatory activity (46, 47).

Another interesting observation of our study was the upregulation of the NAD+ signaling pathway in hyperoxia-exposed PTAFR KO mice compared to WT mice, particularly the increased expression of the mitochondrial enzyme Hmgcs2. Our group has previously shown that mitochondrial bioenergetic and metabolic function of human umbilical venous endothelial cells and mesenchymal stem cells are predictors of risk for BPD in infants (48, 49). Hmgcs2 is a key regulator of ketogenesis which has been found to act as an endogenous protective program that reduces activation of inflammatory macrophages during inflammatory processes that can lead to ARDS (50). Another recent study suggest that Hmgcs2 deficiency may mediate pulmonary fibrosis through alveolar epithelial cell metabolic reprogramming and fibroblast activation (51). Our observation that hyperoxia-exposed PTAFR KO mice had increased expression of SIRT3 which is an important regulator of Hmgcs2 expression in their lungs compared to WT mice suggests that metabolic reprogramming and mitochondrial lipid metabolism may play a role in PAF mediated hyperoxic neonatal lung injury.

A limitation of our study is suggested by our finding that despite better preserved lung structure when exposed to hyperoxia compared to WT mice, PTAFR KO mice had similar lung compliance during hyperoxia. Lung compliance, which is a measure of lung mechanics, is affected by fibrosis and our finding that pathways associated with fibrosis such as the *tumor microenvironment* pathway and the IL-6 related profibrotic *oncostatin-M signaling* pathway are upregulated in both PTAFR KO mice and WT mice is suggestive of a possible mechanistic basis for this observation. A recent study notes that lung compliance measurements in newborn mice exposed to hyperoxia in the first week varies based on the age of the animals suggesting that 14 days (the endpoint used in our study) may be too early a time to detect the improvement in compliance that would be expected based on the improvements in lung structure noted in our study, indicating that additional endpoints such as 4 weeks or 8 weeks might have been useful (52). Next, while hyperoxia-exposed mice are a reproducible model, they are unlikely to replicate all aspects of hyperoxic lung injury or BPD in preterm infants. Another limitation is that we did not measure inflammatory cytokines other than CXCL1 in BAL fluid or cellular infiltrate in the lungs of these mice.

Despite such limitations, the results of our study indicate that PAF signaling may play an important role in promoting inflammation and contribute to impaired alveolar development in hyperoxia-exposed neonatal murine lungs. Future directions could include more detailed analysis of the contribution of the various canonical gene pathways and molecules identified in this study to PAF mediated inflammatory injury in the hyperoxia-exposed newborn lung as well as testing PAF receptor antagonists and PAF acetyl hydrolase in animal models of hyperoxic neonatal lung injury to determine whether PAF signaling could be a useful therapeutic target for the treatment or prevention of BPD.

## ACKNOWLEDGEMENTS

The authors wish to than the expert contributions of Michael R Crowley Ph.D., and David K Crossman Ph.D., UAB Heflin Center Genomics Core. This work was supported by U01 HL122626 (N.A.) and U01 ES02769701 (T.J.).

