## Supplementary material for "Platelet Activating Factor Activity Modulates Hyperoxic Neonatal Lung Injury Severity": Data supplement

**ONLINE DATA SUPPLEMENT**

This appendix has been included to provide readers with additional information about the manuscript **Platelet Activating Factor Activity Modulates Hyperoxic Neonatal Lung Injury Severity**.

**Authors:** Aaron J. Yee, Jegen Kandasamy, Namasivayam Ambalavanan, Changchun Ren, Pankaj Jain, Brian Halloran, Nelida Olave, Teodora Nicola and Tamas Jilling

**SUPPLEMENTAL TABLES**

**Supplemental Table 1:** Top 20 genes up regulated by hyperoxia in C57BL6/J WT mice lungs (genes with > 2 fold change between the 2 groups and FDR P-values < 0.05 in both groups are highlighted in yellow).

| Gene | WT HO vs NO | | PRKO HO vs NO | |
|---|---|---|---|---|
| | Fold Change | FDR P value | Fold Change | FDR P value |
| Ccl2 | 39.49 | 1.22E-17 | 14.00 | 8.82E-19 |
| Eda2r | 32.34 | 2.23E-37 | 24.21 | 7.09E-36 |
| Nppa | 27.79 | 3.22E-04 | 3.01 | 2.20E-01 |
| Chrna7 | 26.58 | 1.20E-10 | 14.62 | 4.57E-10 |
| Saa3 | 23.05 | 2.44E-30 | 12.49 | 5.54E-20 |
| Cxcl10 | 22.34 | 2.09E-18 | 4.33 | 8.97E-06 |
| Tmem255a | 22.26 | 3.84E-35 | 12.94 | 5.91E-26 |
| Cxcl1 | 22.17 | 1.79E-07 | 2.71 | 1.93E-01 |
| 1810006JC | 19.10 | 1.18E-03 | 3.56 | 5.20E-02 |
| Car3 | 17.78 | 1.37E-12 | -1.03 | 9.80E-01 |
| Gdf15 | 17.43 | 2.65E-29 | 14.34 | 2.52E-26 |
| Dio1 | 15.36 | 7.49E-09 | 3.27 | 1.17E-02 |
| Clca3a1 | 14.79 | 9.96E-16 | 24.01 | 8.05E-22 |
| Bpifa1 | 14.36 | 8.22E-08 | 1.50 | 6.54E-01 |
| Fgg | 14.19 | 8.03E-05 | 11.67 | 1.11E-06 |
| Nxpe5 | 13.99 | 1.81E-04 | 3.29 | 2.80E-02 |
| Serpina3i | 13.90 | 6.00E-16 | 9.70 | 1.72E-13 |
| Cdkn1a | 12.46 | 1.08E-54 | 15.64 | 3.25E-65 |
| Apold1 | 11.86 | 7.37E-59 | 6.88 | 8.66E-38 |
| Lif | 11.86 | 2.69E-20 | 5.45 | 1.12E-10 |

**Supplemental Table 2**: Top 20 genes down regulated by hyperoxia in C57BL6/J WT mice lungs (genes with > 2 fold change between the 2 groups and FDR P-values < 0.05 in both groups are highlighted in yellow).

| Gene | WT HO vs NO | | PRKO HO vs NO | |
|---|---|---|---|---|
| | Fold Change | FDR P value | Fold Change | FDR P value |
| Ppy | -19.51 | 2.82E-03 | -84.97 | 6.36E-02 |
| Gm30807 | -20.05 | 3.48E-03 | -8.26 | 5.10E-02 |
| Wfdc8 | -21.47 | 1.02E-04 | -138.29 | 3.22E-02 |
| Ear1 | -24.04 | 1.16E-16 | -8.99 | 1.64E-09 |
| Glb113 | -24.89 | 1.02E-08 | -17.02 | 1.79E-08 |
| Gpat2 | -27.46 | 7.04E-03 | -2.68 | 4.27E-01 |
| Gm17778 | -27.46 | 5.28E-03 | -8.41 | 2.79E-02 |
| Vstm2b | -29.52 | 1.90E-07 | -29.31 | 2.23E-09 |
| Gm40960 | -30.97 | 3.56E-03 | -5.75 | 2.66E-02 |
| Wfdc6b | -33.86 | 9.60E-05 | -34.54 | 1.01E-05 |
| Tmem81 | -37.49 | 4.83E-03 | 1.18 | 9.41E-01 |
| Abca17 | -38.58 | 4.89E-07 | -12.99 | 2.46E-06 |
| Wfdc6a | -43.33 | 5.39E-05 | -8.82 | 3.72E-03 |
| D330041H | -43.60 | 4.00E-07 | -18.90 | 5.69E-05 |
| Ibsp | -43.78 | 1.98E-03 | 1.36 | 8.49E-01 |
| Hist2h3c2 | -59.45 | 5.35E-03 | 1.06 | 9.81E-01 |
| Xlr3b | -69.22 | 3.90E-06 | -1.08 | 9.57E-01 |
| Slc22a8 | -106.17 | 3.48E-02 | -8.18 | 2.25E-01 |
| Hist1h4m | -116.35 | 4.64E-02 | -1.02 | 9.99E-01 |
| Fabp1 | -297.58 | 4.28E-03 | -41.52 | 7.03E-07 |

**Supplemental Table 3:** Top 20 genes up regulated by hyperoxia in PTAFR KO mice lungs (genes with > 2 fold change between the 2 groups and FDR P-values < 0.05 in both groups are highlighted in yellow).

| Gene | WT HO vs NO | | PRKO HO vs NO | |
|---|---|---|---|---|
| | Fold Change | FDR P value | Fold Change | FDR P value |
| LOC108168 | 2.90 | 5.90E-02 | 59.87 | 2.05E-04 |
| Mcpt4 | 5.90 | 6.04E-08 | 25.13 | 2.89E-24 |
| Eda2r | 32.34 | 2.23E-37 | 24.21 | 7.09E-36 |
| Clca3a1 | 14.79 | 9.96E-16 | 24.01 | 8.05E-22 |
| Psrc1 | 11.57 | 1.61E-35 | 21.40 | 1.51E-52 |
| Cma1 | 4.70 | 4.02E-07 | 19.52 | 4.14E-25 |
| Tpsb2 | 4.10 | 2.16E-07 | 17.84 | 1.15E-29 |
| My12 | 1.93 | 5.32E-01 | 16.81 | 2.85E-04 |
| Aldh1a3 | 7.06 | 5.74E-12 | 16.23 | 1.09E-21 |
| Cdkn1a | 12.46 | 1.08E-54 | 15.64 | 3.25E-65 |
| Chrna7 | 26.58 | 1.20E-10 | 14.62 | 4.57E-10 |
| Gdf15 | 17.43 | 2.65E-29 | 14.34 | 2.52E-26 |
| Cacng4 | 2.93 | 1.74E-01 | 14.31 | 1.63E-03 |
| Serpina3n | 9.56 | 4.63E-10 | 14.25 | 9.75E-14 |
| Ccl2 | 39.49 | 1.22E-17 | 14.00 | 8.82E-19 |
| Pla1a | 5.89 | 8.26E-04 | 13.54 | 1.41E-07 |
| Shisa6 | 6.48 | 2.21E-09 | 13.53 | 5.01E-18 |
| Tmem255a | 22.26 | 3.84E-35 | 12.94 | 5.91E-26 |
| Trp53cor1 | 4.63 | 1.06E-03 | 12.91 | 4.20E-07 |
| Ptprn | 7.87 | 5.84E-13 | 12.52 | 2.95E-20 |

**Supplemental Table 4:** Top 20 genes down regulated by hyperoxia in PTAFR KO mice lungs (genes with > 2 fold change between the 2 groups and FDR P-values < 0.05 in both groups are highlighted in yellow).

| Gene | WT HO vs NO | | PRKO HO vs NO | |
|---|---|---|---|---|
| | Fold Change | FDR P value | Fold Change | FDR P value |
| **Kcng4** | -6.54 | 4.58E-03 | -16.98 | 3.56E-03 |
| **Glb113** | -24.89 | 1.02E-08 | -17.02 | 1.79E-08 |
| **Cyp2e1** | -8.55 | 3.36E-11 | -17.12 | 4.53E-19 |
| **Ndst4** | -2.77 | 2.25E-01 | -17.63 | 4.40E-02 |
| **Gm41639** | -7.56 | 4.74E-02 | -17.64 | 4.05E-02 |
| **D330041H** | -43.60 | 4.00E-07 | -18.90 | 5.69E-05 |
| **Zfp663** | -16.73 | 8.00E-11 | -19.69 | 3.10E-11 |
| **Clca1** | -2.03 | 4.72E-01 | -20.60 | 5.45E-05 |
| **Lalba** | -14.00 | 8.97E-03 | -21.33 | 1.88E-02 |
| **Cpa1** | -4.44 | 7.75E-04 | -21.74 | 1.87E-10 |
| **Gm29679** | 4.13 | 3.15E-01 | -21.82 | 1.82E-02 |
| **Adcy8** | -11.86 | 4.25E-22 | -23.34 | 2.49E-34 |
| **Havcr1** | -12.88 | 1.41E-12 | -24.57 | 1.94E-20 |
| **Vstm2b** | -29.52 | 1.90E-07 | -29.31 | 2.23E-09 |
| **Cbln1** | -11.34 | 1.09E-03 | -32.17 | 3.26E-03 |
| **Wfdc6b** | -33.86 | 9.60E-05 | -34.54 | 1.01E-05 |
| **Fabp1** | -297.58 | 4.28E-03 | -41.52 | 7.03E-07 |
| **Crabp1** | -16.55 | 2.56E-20 | -49.59 | 2.73E-36 |
| **Chil4** | -2.04 | 4.53E-01 | -80.28 | 5.74E-10 |
| **Wfdc8** | -21.47 | 1.02E-04 | -138.29 | 3.22E-02 |

**Supplemental Table 5:** Genes included in the *Agranulocyte adhesion and diapedesis* IPA canonical pathway. Genes that were significantly regulated by hyperoxia in the lungs of WT and PTAFR KO mice (fold change values > 2 vs. the other group and P < 0.05 in both groups) are highlighted in yellow.

| Gene | WT HO vs. WT NO | | PRKO HO vs. PRKO NO | |
|---|---|---|---|---|
| | Fold change | FDR p-value | Fold change | FDR p-value |
| **Selp** | 6.759685 | 2.58E-06 | 3.225101 | 0.007481 |
| **Tnf** | 4.456041 | 0.021756 | 1.602264 | 0.583136 |
| **Fn1** | 2.219697 | 4.32E-06 | 2.218993 | 5.57E-06 |
| **Il1r1** | 1.874135 | 6.07E-06 | 2.024292 | 2.56E-07 |
| **Cd34** | 1.675487 | 0.001426 | 1.565714 | 0.010333 |
| **Vcam1** | 1.567566 | 0.004243 | 1.569551 | 0.005855 |
| **Glg1** | 1.312161 | 0.384348 | 1.146323 | 0.767979 |
| **Aoc3** | 1.261207 | 0.32016 | -1.2348 | 0.459018 |
| **Tnfrsf1a** | 1.234967 | 0.213738 | 1.293754 | 0.154684 |
| **Itgb2** | 1.181148 | 0.601407 | 1.382972 | 0.312213 |
| **Itga4** | 1.169271 | 0.658067 | -1.20085 | 0.675 |
| **Sdc4** | 1.142222 | 0.517468 | 1.252041 | 0.288019 |
| **C5ar1** | -1.00462 | 0.99182 | 1.099649 | 0.848105 |
| **Cdh5** | -1.00774 | 0.978549 | -1.02123 | 0.956796 |
| **Selplg** | -1.01598 | 0.964504 | -1.11606 | 0.781966 |
| **Itgb7** | -1.08086 | 0.812153 | -1.40648 | 0.229674 |
| **Pecam1** | -1.10581 | 0.680966 | -1.14574 | 0.638465 |
| **Itga5** | -1.25895 | 0.189483 | -1.16987 | 0.490854 |
| **Jam3** | -1.26349 | 0.25703 | -1.25132 | 0.358253 |
| **Cxcl12** | -1.75117 | 0.000254 | -2.41075 | 1E-09 |
| **Sell** | -1.83618 | 0.130091 | -3.17194 | 0.001571 |
| **Cxcr4** | -7.25694 | 9.46E-39 | -5.90032 | 3.06E-31 |

**Supplemental Table 6:** Genes included in the *Tumor microenvironment* IPA canonical pathway. Genes that were significantly upregulated by hyperoxia in the lungs of WT and PTAFR KO mice (fold change values > 2 vs. the other group and P < 0.05 in both groups) are highlighted in yellow (WT higher than PTAFR KO) or blue (PTAFR KO higher than WT).

| Gene | WT HO vs. WT NO | | PRKO HO vs. PRKO NO | |
|---|---|---|---|---|
| | **Fold change** | **FDR p-value** | **Fold change** | **FDR p-value** |
| **IL6** | 59.13956 | 0.000334 | 18.35983 | 0.000402 |
| **Ccl2** | 39.48902 | 1.22E-17 | 13.99874 | 8.82E-19 |
| **Tnf** | 4.456041 | 0.021756 | 1.602264 | 0.583136 |
| **Ptgs2** | 4.301681 | 2.07E-06 | 1.871583 | 0.103458 |
| **Plau** | 4.051229 | 1.08E-17 | 3.606383 | 1.73E-15 |
| **Spp1** | 4.00319 | 2.08E-07 | 8.178234 | 3.16E-17 |
| **Ccnd1** | 3.272113 | 4.23E-16 | 2.93512 | 2.72E-13 |
| **Myc** | 2.7466 | 2.46E-07 | 1.906007 | 0.002756 |
| **Icam1** | 2.670234 | 2.38E-06 | 2.442992 | 3.05E-05 |
| **Cd274** | 2.239314 | 0.0039 | 1.603665 | 0.165551 |
| **Fn1** | 2.219697 | 4.32E-06 | 2.218993 | 5.57E-06 |
| **Mmp2** | 1.965964 | 7.86E-05 | 2.159559 | 5.68E-06 |
| **Hif1a** | 1.937688 | 3.43E-06 | 1.638422 | 0.001338 |
| **Cspg4** | 1.929301 | 6.65E-06 | 1.772906 | 0.000156 |
| **Osm** | 1.874914 | 0.182695 | 1.108229 | 0.899694 |
| **Tnc** | 1.719295 | 0.095007 | 1.992335 | 0.036999 |
| **Il1b** | 1.601683 | 0.325778 | -1.59882 | 0.419898 |
| **Mmp9** | 1.532594 | 0.322112 | -1.2173 | 0.765719 |
| **Stat3** | 1.447066 | 0.015706 | 1.593646 | 0.002018 |
| **Slc16a1** | 1.424905 | 0.072659 | 1.257837 | 0.364202 |
| **Csf1** | 1.322545 | 0.071239 | 1.219634 | 0.298616 |
| **Cd44** | 1.295075 | 0.117707 | 1.386821 | 0.054065 |
| **Jak2** | 1.287034 | 0.190664 | 1.193695 | 0.475608 |
| **Bcl2** | 1.286652 | 0.14705 | 1.143048 | 0.58229 |
| **Hgf** | 1.257147 | 0.614868 | 1.139755 | 0.839728 |
| **Tnfrsf1a** | 1.234967 | 0.213738 | 1.293754 | 0.154684 |
| **Tslp** | 1.230161 | 0.56992 | -1.27743 | 0.572146 |
| **Nos2** | 1.220748 | 0.431322 | -1.06899 | 0.866125 |
| **Map3k14** | 1.214431 | 0.373946 | 1.032904 | 0.932652 |
| **Rac1** | -1.00836 | 0.975304 | -1.08609 | 0.780699 |
| **Tiam1** | -1.06234 | 0.803117 | -1.46091 | 0.038649 |
| **Csf2** | -1.06457 | 0.915171 | 1.077899 | 0.92644 |

| | | | | |
|---|---|---|---|---|
| **Fas** | -1.07519 | 0.7666 | -1.04474 | 0.899694 |
| **Arg1** | -1.1679 | 0.804246 | -1.08685 | 0.927681 |
| **Bad** | -1.24866 | 0.256811 | 1.014864 | 0.969784 |
| **Slc1a4** | -1.29363 | 0.12107 | -1.54198 | 0.005306 |
| **Cflar** | -1.40107 | 0.043851 | -1.22344 | 0.354672 |
| **Cxcl12** | -1.75117 | 0.000254 | -2.41075 | 1E-09 |
| **Lepr** | -2.05981 | 0.000632 | -2.8598 | 1.8E-07 |
| **Cxcr4** | -7.25694 | 9.46E-39 | -5.90032 | 3.06E-31 |

**Supplemental Table 7:** Canonical pathways regulated with Z(abs) ≥ 1 and -logP >1.3 (P<0.05).

by hyperoxia in WT mice, PTAFR KO mice and both groups.

| Group | Canonical Pathway | -logP | Ratio | Z-score |
|---|---|---|---|---|
| WT | Role of Hypercytokinemia/hyperchemokinemia in the Pathogenesis of Influenza | 3.45 | 0.116 | 3.162 |
| | Tumor Microenvironment Pathway | 2.36 | 0.0726 | 2.496 |
| | Xenobiotic Metabolism CAR Signaling Pathway | 2.15 | 0.0684 | 2.496 |
| | p38 MAPK Signaling | 1.45 | 0.0667 | 2.121 |
| | IL-17 Signaling | 3.55 | 0.0856 | 2 |
| | Differential Regulation of Cytokine Production in Intestinal Epithelial Cells by IL-17A and IL-17F | 2.28 | 0.174 | 2 |
| | Interferon Signaling | 1.59 | 0.111 | 2 |
| | LXR/RXR Activation | 2.77 | 0.0894 | -2.333 |
| Both | CREB Signaling in Neurons | 6.84 | 0.0693 | -2.655 |
| | Oncostatin M Signaling | 2.13 | 0.116 | 2.236 |
| | Tumor Microenvironment Pathway | 4.79 | 0.095 | 2.183 |
| | Sperm Motility | 1.48 | 0.051 | -2 |
| | Adrenomedullin signaling pathway | 2.3 | 0.0653 | -2.496 |
| | Opioid Signaling Pathway | 1.81 | 0.0536 | -2.496 |
| | p70S6K Signaling | 1.44 | 0.0606 | -2.646 |
| PTAFR-KO | NAD Signaling Pathway | 1.45 | 0.0331 | 2.236 |
| | IL-13 Signaling Pathway | 1.89 | 0.0431 | -2.236 |

#### SUPPLEMENTAL FIGURE LEGENDS

**Supplemental Figure 1**: **Pathway map of the *NAD signaling* pathway**. Known interactions and interrelationships between the component genes of the *NAD signaling* pathway that was identified to be differentially regulated by hyperoxia in the lungs of PTAFR KO mice only and not in the lungs of WT mice.

**Supplemental Figure 2**: **HMGCS2 and SIRT3 expression in the lung**. (**A**) HMGCS2 and (**B**) SIRT3 expression measured using RNA isolated from lung tissue analyzed using real time polymerase chain reaction (RT-PCR), two-way ANOVA and Tukey test for post hoc analysis.

**Supplemental Figure 3**: **Pathway map of the *tumor microenvironment* pathway**. Known interactions and interrelationships between the component genes of the *tumor microenvironment* pathway that was identified to be differentially regulated by hyperoxia in the lungs of both WT and PTAFR KO mice.

### Supplemental Figure 1

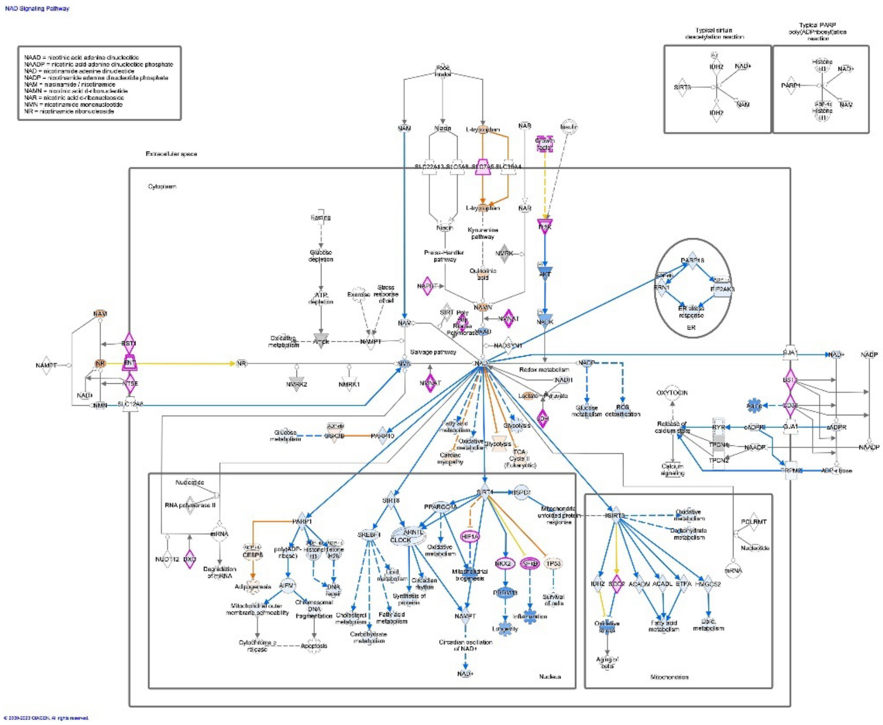


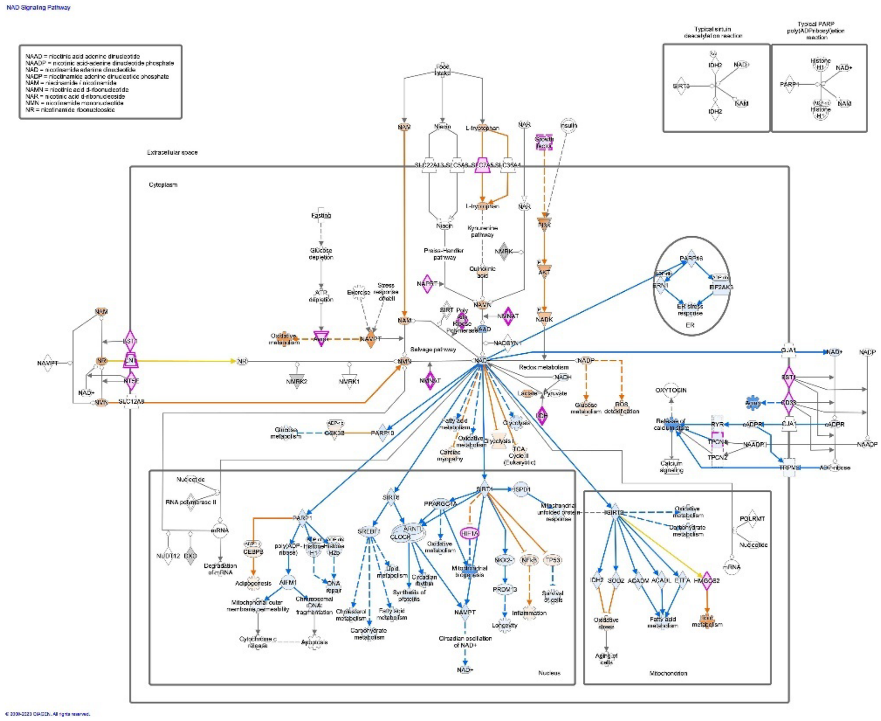

**Supplemental Figure 2**

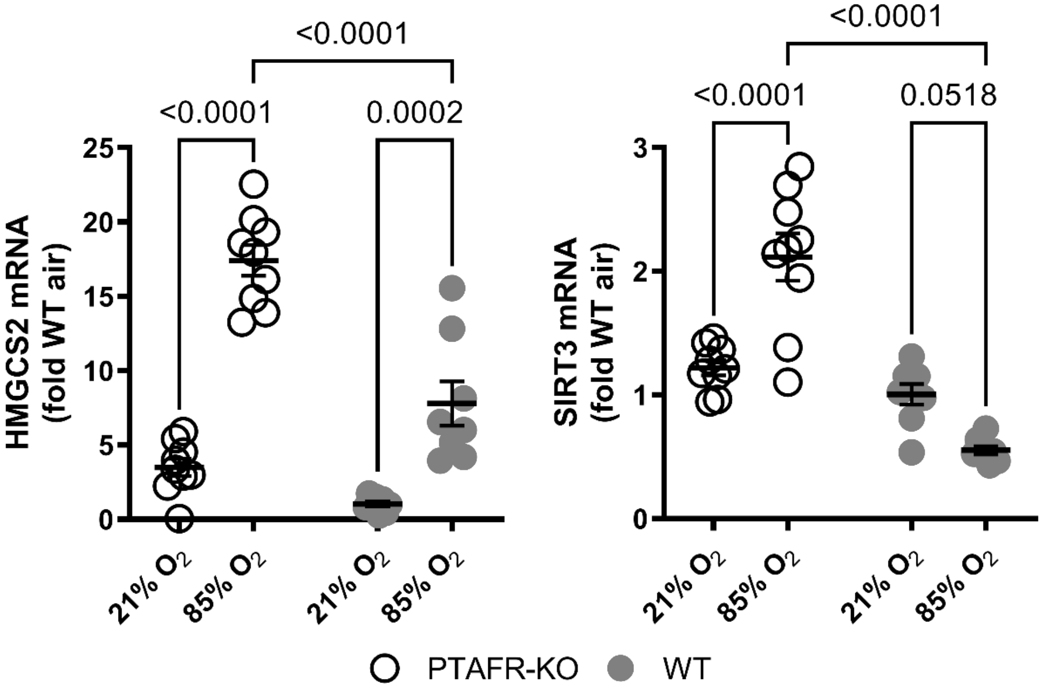

### Supplemental Figure 3

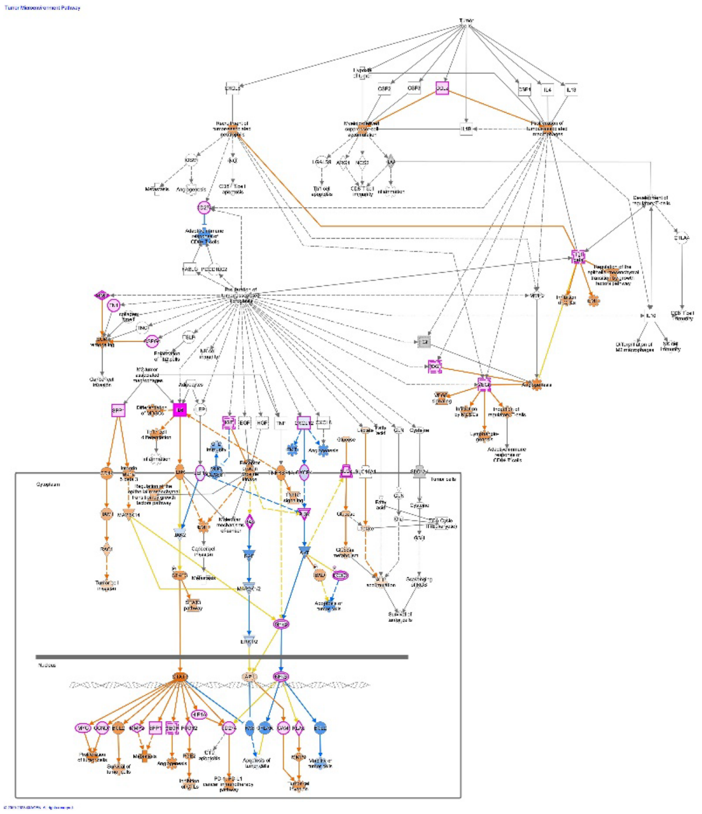

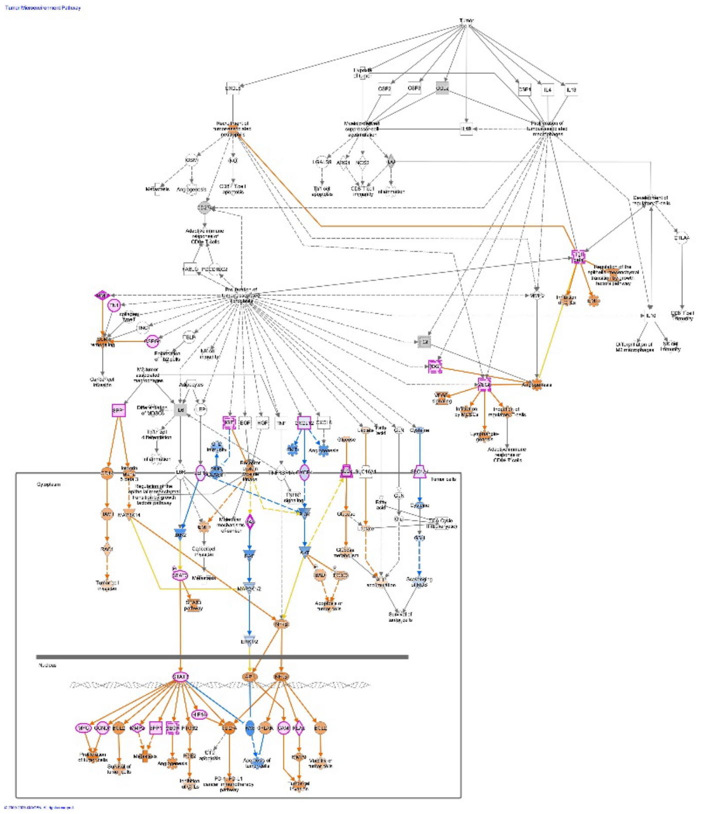
